# Delta breakthrough infections elicit potent, broad and durable neutralizing antibody responses

**DOI:** 10.1101/2021.12.08.471707

**Authors:** Alexandra C. Walls, Kaitlin R. Sprouse, Anshu Joshi, John E. Bowen, Nicholas Franko, Mary Jane Navarro, Cameron Stewart, Matthew McCallum, Erin A. Goecker, Emily J. Degli-Angeli, Jenni Logue, Alex Greninger, Helen Chu, David Veesler

**Affiliations:** Department of Biochemistry, University of Washington, Seattle, WA 98195, USA; Howard Hughes Medical Institute, University of Washington, Seattle, WA 98195, USA; Division of Allergy and Infectious Diseases, University of Washington, Seattle, WA 98195, USA; Department of Laboratory Medicine and Pathology, University of Washington School of Medicine, Seattle, WA, USA

## Abstract

The SARS-CoV-2 Delta variant is currently responsible for most infections worldwide, including among fully vaccinated individuals. Although these latter infections are associated with milder COVID-19 disease relative to unvaccinated subjects, the specificity and durability of antibody responses elicited by Delta breakthrough cases remain unknown. Here, we demonstrate that breakthrough infections induce serum binding and neutralizing antibody responses that are markedly more potent, durable and resilient to spike mutations observed in variants of concern than those observed in subjects who were infected only or received only two doses of COVID-19 vaccine. However, wee show that Delta breakthrough cases, subjects who were vaccinated after SARS-CoV-2 infection and individuals vaccinated three times (without infection) have serum neutralizing activity of comparable magnitude and breadth indicate that multiple types of exposure or increased number of exposures to SARS-CoV-2 antigen(s) enhance spike-specific antibody responses. Neutralization of the genetically divergent SARS-CoV, however, was moderate with all four cohorts examined, except after four exposures to the SARS-CoV-2 spike, underscoring the importance of developing vaccines eliciting broad sarbecovirus immunity for pandemic preparedness.

---

The SARS-CoV-2 Delta (B.1.617.2) variant of concern emerged at the end of 2020 and became dominant globally by mid-2021. Mutations in the spike (S) glycoprotein (Johnson et al., 2021; Walls et al., 2020a; Wrapp et al., 2020) and in the nucleoprotein (N) have been suggested to account for its enhanced transmissibility, replication kinetics and viral loads in oropharyngeal and nose-throat swabs of infected individuals relative to the ancestral Wuhan-Hu-1 virus and other variants (Li et al., 2021; Liu et al., 2021; Mlcochova et al., 2021; Saito et al., 2021; Syed et al., 2021). Moreover, multiple S mutations in the N-terminal domain and receptor-binding domain have been shown to promote immune evasion (McCallum et al., 2021a, 2021b; Mlcochova et al., 2021; Suryadevara et al., 2021; Ying et al., 2021). These characteristics combined with waning of serum neutralizing antibody titers over time in vaccinated individuals has resulted in Delta breakthrough infections that are usually associated with much milder symptoms than infection of unvaccinated individuals (Levine-Tiefenbrun et al., 2021; Mlcochova et al., 2021).

Understanding the magnitude and breadth of immune responses following a breakthrough infection is key to guiding vaccination policies and pandemic preparedness efforts (Collier et al., 2021). Serum neutralizing antibody titers represent the current best correlate of protection against SARS-CoV-2 in animal challenge studies (Arunachalam et al., 2021; Case et al., 2020a; Corbett et al., 2021; Hassan et al., 2021; Khoury et al., 2021; McMahan et al., 2021; Winkler et al., 2020) and multiple clinical trials have shown the benefits of therapeutic administration of monoclonal antibodies in humans (Corti et al., 2021). Furthermore, serum neutralizing antibodies are used in ongoing comparative clinical trials as key success metrics for the next generation of vaccines (e.g., NCT05007951 and NCT04864561 comparing GBP510 and VLA2001 to AZD1222, respectively). To understand whether the order of infection and/or vaccination as well as repeated exposures alter the specificity, magnitude, and breadth of antibody responses, we followed and compared serum antibodies in individuals who were vaccinated, who were previously infected and then vaccinated, or who were first vaccinated and then infected with the SARS-CoV-2 Delta variant.

We compared serum binding titers following infection, vaccination, or both in groups of ∼15 individuals enrolled through the longitudinal cohort study, HAARVI, at the University of Washington in Seattle (**Table S1-S4)**. Individuals in the Delta breakthrough group (n=1 with Johnson and Johnson Ad26.COV2.S, n=2 with Moderna mRNA-1273, n=13 with Pfizer Cominarty), in the infected then vaccinated (infected/vaccinated) cohort (n=1 with Johnson and Johnson Ad26.COV2.S, n=3 with Moderna mRNA-1273, n=11 with Pfizer Cominarty), and in the vaccinated-only group (n=3 with Moderna mRNA-1273, n=12 with Pfizer Cominarty) (**Table S1-S3**) were compared to human convalescent sera (HCS) which were collected prior to October 2020 in Washington State (all samples were obtained prior to July 2020 except one which was drawn in September 2020), indicating these infections were not with any variants of concern (VOC) (according to outbreaks.info) (**Table S4**). Eight individuals from the infected/vaccinated or vaccinated-only groups received a third vaccine dose (i.e. booster, designated 3X). All these samples were compared to SARS-CoV-2 naive individuals who had blood drawn prior to vaccination (**Table S5**) as confirmed by the lack of SARS-CoV-2 nucleocapsid (N) reactivity using the Roche Elecsys anti-N immuno assay (only convalescent samples were positive) (**Table S1-S5**).

Serum IgG, IgA, and IgM binding titers were evaluated using ELISAs with the SARS-CoV-2 Hexapro S antigen (Hsieh et al., 2020). The cohorts were followed longitudinally for up to 6 months/180 days after initial blood draws to evaluate differences in durability of antibody responses. Responses were highest amongst individuals who were exposed to SARS-CoV-2 S three or four times through vaccination-only or a combination of infection and vaccination. The magnitude of IgG responses for vaccinated individuals who experienced a breakthrough infection with the Delta variant had geometric mean titers (GMT) of 2.1×10^5^ and 8.8×10^4^ 30 days and 60 days after positive PCR test, respectively (**Figure 1A and Figure S1**). Infected/vaccinated individuals had binding GMTs of 1.3×10^5^, 5.8×10^4^ and 7.8×10^4^ at 10 days, 112 days and 180 days after the second dose and 4.6×10^5^ 10 days after a third vaccine shot (**Figure 1A and Figure S1**), corresponding to a 3.6 fold increase when comparing titers 10 days after the second and third vaccination. Vaccinated-only individuals had binding GMTs of 1.7×10^4^, 9.5×10^3^ and 5.4×10^3^ at 10 days, 112 days and 180 days after receiving a second vaccine dose which rose to 4.6×10^5^ 10 days following a boost with a third vaccination (**Figure 1A and Figure S1**), amounting to a 26 fold increase between the peak titers after the second and third vaccination. We determined GMTs of 4.2×10^2^ for infected individuals that were not vaccinated and 8.8×10^1^ (just above the limit of detection) for SARS-CoV-2 naive individuals, as confirmed by the lack of reactivity in a SARS-CoV-2 N immunoassay (**Figure 1A, Figure S1, Table S5)**. These data suggest that repeated exposures through vaccination, infection or a combination of both induce potent and durable polyclonal plasma antibody binding titers. Furthermore, we observed a greater durability of binding GMTs over 180 days for the infected/vaccinated relative to the vaccinated-only (two doses) cohorts, which could result from increased number of exposures or due to actual infection.

**Figure 1:**
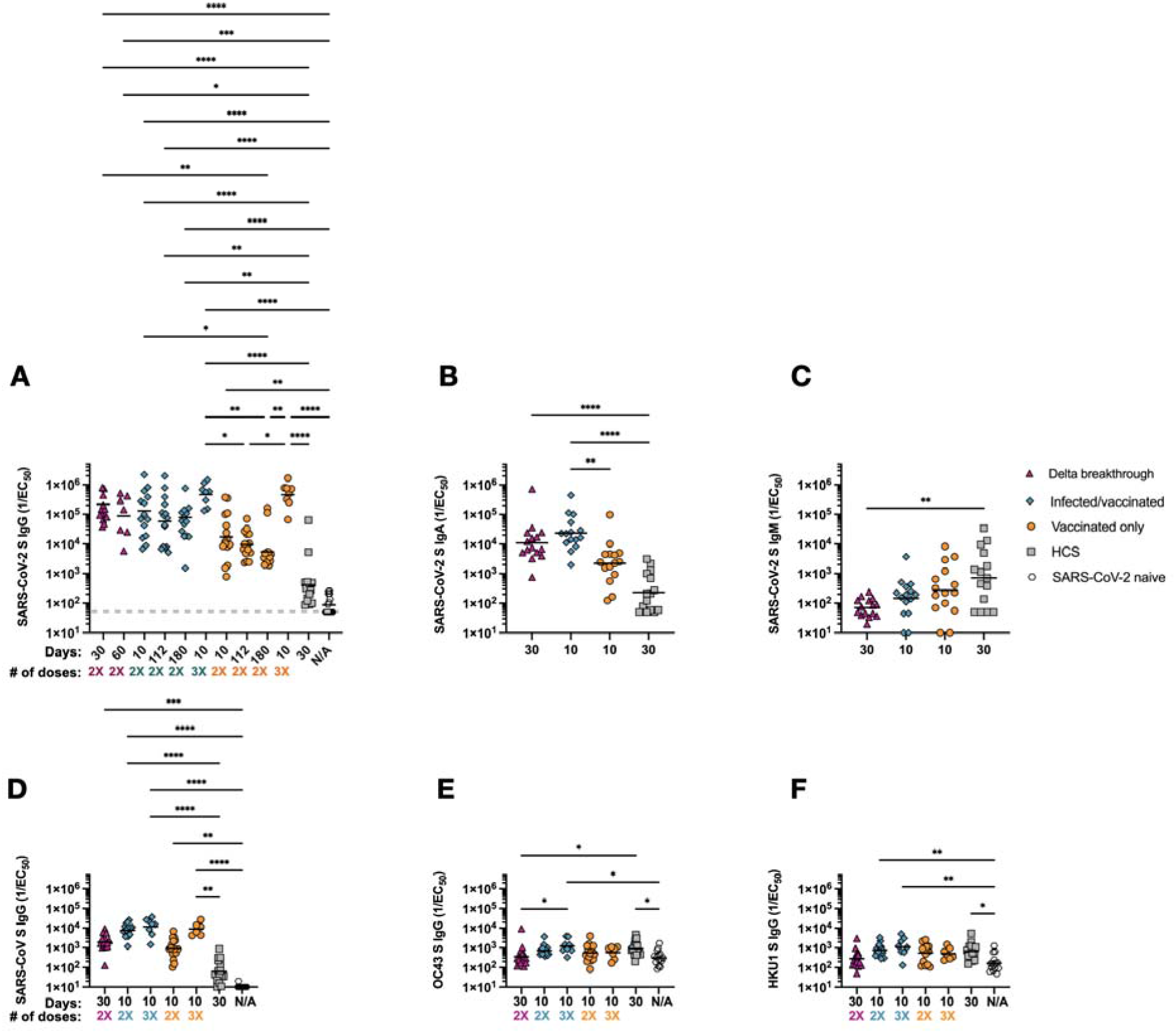
Repeated exposures to SARS-CoV-2 antigens through vaccination or infection enhance S-specific serum IgG and IgA binding titers. **(A)** Serum IgG binding titers at 30 or 60 days post infection or 10, 112, or 180 days post second or third vaccine dose were evaluated for longitudinal samples by ELISA using prefusion-stabilized SARS-CoV-2 S Hexapro as antigen. Serum samples were obtained from individuals who had a Delta breakthrough infection (n=15, magenta triangle), were previously infected then vaccinated (n=15, teal diamond), have only been vaccinated (n=15, orange circle), were infected in 2020 in Washington State (n=15, gray square), or were SARS-CoV-2 naive (samples taken prior to vaccination, n=15, open hexagon). **(B)** Serum IgA binding titers at 30 days post infection or 10 days post second vaccine dose were evaluated by ELISA using prefusion-stabilized SARS-CoV-2 S Hexapro as antigen. **(C)** Serum IgM binding titers at 30 days post infection or 10 days post second vaccine dose were evaluated by ELISA using prefusion-stabilized SARS-CoV-2 S Hexapro as antigen. **(D)** Serum IgG binding titers were evaluated by ELISA at 30 days post infection, 10 days post second or third vaccine dose or prior to SARS-CoV-2 exposure (SARS-CoV-2 naive) using prefusion-stabilized SARS-CoV 2P S as antigen. **(E-F)** Serum IgG binding titers were evaluated at 30 days post infection, 10 days post second or third vaccine dose, or prior to SARS-CoV-2 exposure (SARS-CoV-2 naive) by ELISA using OC43 S (E) or HKU1 2P S (F) as antigen. # of doses: number of vaccine doses received. Statistical significance was determined by Kruskal Wallis and Dunn’s multiple comparisons test and shown only when significant. \**P* < 0.05; \*\**P* < 0.01; \*\*\**P* < 0.001; and ****P < 0.0001. LOD is shown as a gray horizontal dotted line when above the x axis. Raw fits are shown in Figure S1 and S2.

We determined peak S-specific IgA binding GMTs of 1.1×10^4^ for Delta breakthrough cases, 2.3×10^4^ for infected/vaccinated individuals, 2.2×10^3^ for vaccinated-only individuals and 2.3×10^2^ for HCS (**Figure 1B and Figure S2)**, following a magnitude trend similar to IgG binding titers. Conversely, IgM binding titers were roughly an order of magnitude higher for HCS (GMT of 7×10^2^) relative to other cohorts (**Figure 1C)**, presumably as a result of extensive immunoglobulin class switching and affinity maturation occurring for the latter cases, mirroring findings made with memory B cells in vaccinated individuals (Cho et al., 2021). Collectively, these data indicate that breakthrough infections, vaccination following infection and triple vaccination trigger similarly robust anamnestic responses.

To assess how vaccination and infection affect the breadth of binding antibody responses, we performed ELISAs using prefusion diproline-stabilized (2P) SARS-CoV S as antigen (Kirchdoerfer et al., 2018; Pallesen et al., 2017). At 30 days post infection or 10 days post second vaccination (2X), we determined GMTs of 1.9×10^3^ for Delta breakthrough patients, 7.7×10^3^ for infected/vaccinated individuals, and 9.3×10^2^ for vaccinated-only individuals (**Figure 1D**). Upon receiving a booster immunization, both infected/vaccinated (3X) and vaccinated-only (3X) SARS-CoV S-specific responses increased to 1.2×10^4^ (1.5 fold increase from 2X) and 8.9×10^3^ (9.5 fold increase from 2X), respectively. We observed the lowest SARS-CoV S responses for HCS (6.3×10^1^), which are subjects who had only been infected once by SARS-CoV-2, and for naive individuals (limit of detection), who were never exposed. These results suggest that antibody cross-reactivity within the sarbecovirus subgenus improves markedly with repeated exposures to SARS-CoV-2 S (**Figure 1D)**. We also investigated antibody cross-reactivity beyond the sarbecovirus subgenus using the prefusion-stabilized S trimers of the common-cold causing embecoviruses OC43 (Tortorici et al., 2019) and HKU1 as ELISA antigens. We determined GMTs of 3.6×10^2^ (OC43 S) and 2.8×10^2^ (HKU1 S) for Delta breakthrough, 7.2×10^2^ (OC43 S) and 7.6×10^2^ (HKU1 S) for 2X infected/vaccinated, 1.2×10^3^ (OC43 S) and 1.1×10^3^ (HKU1 S) for 3X infected/vaccinated, 5.7×10^2^ (OC43 S) and 5.5×10^2^ (HKU1 S) for 2X vaccinated-only, and 5.8×10^2^ (OC43 S) and 5.1×10^2^ (HKU1 S) for 3X vaccinated-only cohorts (**Figure 1E-F**). Individuals who were infected with SARS-CoV-2 had GMTs of 9.2×10^2^ (OC43 S) and 6.7×10^2^ (HKU1 S) and those who were SARS-CoV-2 naive had GMTs of 3.1×10^2^ (OC43 S) and 1.7×10^2^ (HKU1 S) (**Figure 1E-F**). These findings show that exposure to SARS-CoV-2 S does not enhance emebecovirus cross-reactive antibody responses even compared to SARS-CoV-2 naive individuals, who might have been previously exposed to seasonal coronaviruses.

To understand the magnitude and durability of neutralizing antibody responses among the different groups, we determined serum neutralizing activity for all samples obtained longitudinally using vesicular stomatitis virus (VSV) pseudotyped with SARS-CoV-2 G614 S (Kaname et al., 2010; Lempp et al., 2021). Delta breakthrough infections resulted in neutralizing GMTs of 2.7×10^3^ after 30 days and 2.9×10^3^ after 60 days (**Figure 2A, Figures S3-S4)**. We also observed potent neutralizing activity for infected/vaccinated subject sera with GMTs of 3.1×10^3^, 2.6×10^3^ and 1.8×10^3^ at 10 days, 112 days and 180 days post second vaccine dose, respectively, and increased 2.5 fold to 7.6×10^3^ 10 days post third vaccine dose (**Figure 2A, Figures S5-S8)**. Vaccinated-only individuals had serum neutralizing activity of 3.7×10^2^, 1.4×10^2^ and 1.3×10^2^ at 10, 112 and 180 days post second vaccine dose which rose to 3.1×10^3^ after a third vaccine dose (**Figure 2A, Figures S9-S12)**. This corresponds to an 8.2 fold increase when comparing 10 days after each dose, yielding a potency comparable to Delta breakthrough cases 30 days after infection and infected/vaccinated subjects 10 days post second vaccine dose. HCS had weak neutralizing potency (GMT 3.5×10^1^) (**Figure 2A, Figure S13**). These data suggest that the number of exposures to S is more important than the type of exposure (infection vs. vaccination) to determine the magnitude of serum neutralizing activity. However, the approximately ten fold greater and constant neutralizing antibody GMTs measured over time for Delta breakthrough and infected/vaccinated cohorts set these 2 groups apart compared to vaccinated-only individuals (who experienced a neutralizing GMT decay of ∼55% over 6 months after the second vaccine dose). These findings are in agreement with studies of individuals infected and subsequently vaccinated with mRNA vaccines or Jansen Ad26.COV2.S (Keeton et al., 2021; Krammer et al., 2021; Saadat et al., 2021; Stamatatos et al., 2021). Hence, multiple types of exposure or increased number of exposures to SARS-CoV-2 antigen(s) enhance the magnitude and durability of serum neutralizing activity possibly due to sustained antibody production by long lived plasma cells, increased affinity maturation yielding antibodies with greater neutralization potency, or a combination of both.

**Figure 2:**
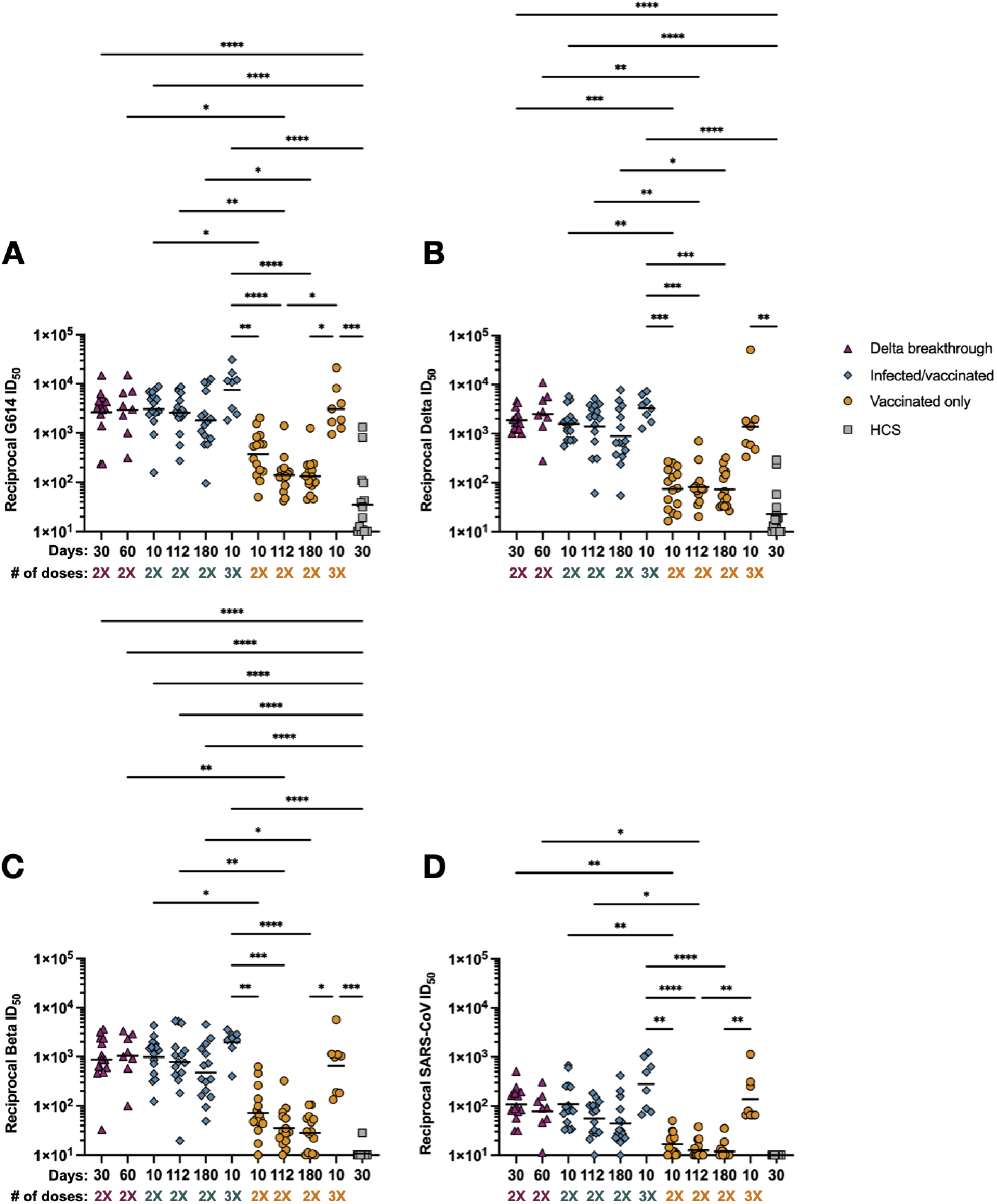
Delta breakthrough, infected/vaccinated and triple vaccinated individuals have exceptionally high serum neutralizing activity. Serum samples were obtained from individuals who had a Delta breakthrough infection (n=14, magenta triangle), who were previously infected then vaccinated (n=15, teal diamond, infected/vaccinated), who have been vaccinated only (n=15, orange circle), or who were infected only in 2020 in Washington State (n=15, gray square, HCS). All neutralization assays were performed using VeroE6-TMPRSS2 as target cells at least in duplicate. (**A**) SARS-CoV-2 G614 S VSV pseudotype neutralization. (**B**) SARS-CoV-2 Delta S VSV pseudotype neutralization. (**C**) SARS-CoV-2 Beta S VSV pseudotype neutralization. (**D**) SARS-CoV S VSV pseudotype neutralization. # of doses: number of vaccine doses received. Statistical significance was determined by Kruskal Wallis and Dunn’s multiple comparisons test and shown only within group or at matched timepoint for ease of viewing when significant. \**P* < 0.05; \*\**P* < 0.01; \*\*\**P* < 0.001; and ****P < 0.0001. Normalized curves and fits are shown in Fig. S3-S13.

We also observed durable neutralizing activity for the Delta breakthrough serum samples against both Delta S and Beta S VSV pseudoviruses for which we determined GMTs of 1.9×10^3^ (Delta S) and 8.8×10^2^ (Beta S) after 30 days and 2.5×10^3^ (Delta S) and 1.0×10^3^ (Beta S) after 60 days **(Figure 2B-C)**. The dampening in neutralization potency relative to G614 S VSV for both Delta S (1.2-1.4 fold GMT reduction) and Beta S (2.9-3.0 fold GMT reduction) was smaller than previous findings with vaccinated-only or convalescent cohorts (Cele et al., 2021; Madhi et al., 2021; McCallum et al., 2021a; Planas et al., 2021; Walls et al., 2021a; Wibmer et al., 2021). For infected/vaccinated subjects, we determined GMTs of 1.6×10^3^, 1.4×10^3^, and 8.9×10^2^ at 10, 112 and 180 days post second vaccine dose against Delta S VSV, respectively, corresponding to a 1.8-2.0 fold GMT reduction relative to G614 S VSV **(Figure 2A-B)**. After receiving a third vaccine dose, these individuals experienced a 2.0 fold increase in neutralizing titers against Delta S VSV (GMT 3.3×10^3^), corresponding to a 2.3 fold GMT reduction relative to G614 S VSV. Therefore, although the magnitude of neutralizing GMT increased, the breadth of broadly neutralizing antibody responses remains similar against Delta S VSV **(Figure 2A-B)**. Beta S VSV neutralizing GMTs were 9.9×10^2^, 7.9×10^2^ and 4.8×10^2^ at 10, 112 and 180 days post second vaccine dose, respectively, corresponding to a 3.1-3.8 GMT fold change compared to G614 S VSV **(Figure 2A,C)**. Following the third vaccination, these individuals experienced a 2.0 fold increase in neutralizing titers against Beta S VSV (GMT 1.9×10^3^), yielding a 3.9 fold GMT reduction relative to G614 S VSV. The vaccinated-only cohort had neutralizing GMTs against Delta S VSV of 7.5×10^1^, 8.2×10^1^ and 7.4×10^1^ at 10, 112 and 180 days post second vaccine dose, respectively, corresponding to a 1.7-5 fold GMT reduction relative to G614 S VSV (**Figure 2B)**. Triple vaccinated-only subjects had neutralizing antibody GMTs of 1.4×10^3^, an 18.8 fold increase compared to the same individuals after receiving only two vaccine doses at the same time point (**Figure 2B)**. Beta S VSV pseudovirus neutralizing GMTs were 7.3×10^1^, 3.6×10^1^ and 2.8×10^1^ at 10, 112 and 180 days post second vaccine dose, respectively (**Figure 2C)**. This corresponds to a 4.0-5.1 fold GMT reduction of serum neutralizing activity relative to G614 S VSV. Triple vaccinated-only subjects displayed a 9.0 fold neutralizing GMT increase (6.6×10^2^) compared to the same time point after receiving only two vaccine doses, consistent with enhanced resilience to this variant. These data suggest that repeated exposures, even to a distinct antigen, improves the potency and resilience of serum neutralizing activity to variants, including against Beta S VSV (Walls et al., 2021b) for which an average ∼7 fold dampening of potency has been reported in the literature (mostly comprising infected-only and vaccinated-only subjects) (https://covdb.stanford.edu/), in line with previous studies of infected/vaccinated individuals (Keeton et al., 2021; Stamatatos et al., 2021). Moreover, we observed a subtle advantage of mixed cohorts, including subjects who experienced both infections and vaccination (irrespective of the order) in terms of neutralization breadth against SARS-CoV-2 variants relative to triple vaccinated-only individuals. It remains to be determined if waning of binding and neutralizing GMTs will occur with similar kinetics after the third immunization of vaccinated-only subjects than after two doses or if they will mirror the durable antibody responses observed in Delta breakthrough and infected/vaccinated cohorts.

To assess the breadth of serum neutralizing activity among these different cohorts against viruses belonging to a different sarbecovirus clade than SARS-CoV-2, we used a SARS-CoV S VSV pseudotype. Delta breakthrough cases had neutralizing GMTs of 1.1×10^2^, and 7.9×10^1^ at 30 and 60 days post positive PCR test (**Figure 2D**) whereas the 2X infected/vaccinated cohort had GMTs of 1.1×10^2^, 5.6×10^1^ and 4.4×10^1^ at 10, 112 and 180 days post second vaccine dose, respectively (**Figure 2D**). This corresponds to GMT reductions of 24-36-fold (Delta breakthrough) and 28-46 fold (infected/vaccinated) compared to G614 S VSV. After receiving a third vaccine dose, infected/vaccinated individuals had ∼2.5 fold enhanced serum neutralizing responses against SARS-CoV S VSV (GMT of 2.8×10^2^) relative to the same subjects at 10 days post second vaccination. Although this amounts to a 26 fold reduction compared to peak neutralizing G614 S VSV GMT,, this is a similar neutralizing activity than that of vaccinated-only individuals against G614 after 2 doses (GMT 3.7×10^2^). The vaccinated-only cohort had neutralizing GMTs of 1.7×10^1^, 1.3×10^1^ and 1.2×10^1^ at 10, 112 and 180 days post second vaccine dose, respectively, which were boosted to 1.4×10^2^ after the third vaccination (**Figure 2D**). The latter GMT matches serum neutralizing activity of Delta breakthrough cases or of infected/vaccinated individuals (after two vaccinations),, further suggesting that the number of exposures is more important than the type of exposure. No SARS-CoV neutralization was detected for HCS, in line with the weak overall serum neutralizing activity of this cohort across any sarbecoviruses (**Figure 2D**).

We show that Delta breakthrough cases or infected/vaccinated subjects are endowed with greater potency, breadth and durability of serum neutralizing activity relative to individuals who received 2 doses of COVID-19 vaccine or those who were only infected by SARS-CoV-2 in 2020. Serum neutralizing GMTs for the Delta breakthrough cases and infected/vaccinated subjects were greater against the Delta and Beta S variants at all time points than that of vaccinated-only individuals (after two doses) against G614 S pseudovirus at peak titers. The latter levels of neutralization potency were associated with ∼95% protection from symptomatic COVID-19 disease in phase 3 clinical trials of the Pfizer Cominarty (Polack et al., 2020) and Moderna mRNA-1273 (Baden et al., 2021) vaccines and are reminiscent of observations made with infected/vaccinated samples (Stamatatos et al., 2021). The marked enhancement of serum neutralizing activity for vaccinated-only subjects after three doses, to levels comparable to Delta breakthrough and infected/vaccinated cases, suggest that the number of exposures to SARS-CoV-2 S, whether through vaccination or infection, correlates with the strength of neutralizing antibody responses as well as resilience to variants. It remains to be determined whether the number of exposures also correlates with durability of antibody responses for vaccinated-only individuals, as shown here with Delta breakthrough cases up to 60 days and infected/vaccinated subjects for 180 days. The reduction of serum neutralizing activity against SARS-CoV presented here following up to 3 SARS-CoV-2 S exposures, along with previous studies (Martinez et al., 2021; Stamatatos et al., 2021; Walls et al., 2021a), suggest that combinations of COVID-19 disease and vaccination would leave the population vulnerable to infection by a genetically divergent sarbecovirus, due to the more extensive sequence differences between SARS-CoV-2 S and SARS-CoV S compared to any two SARS-CoV-2 S variants. Four exposures to SARS-CoV-2 S, through infection followed by three vaccinations, elicit SARS-CoV serum neutralizing titers with a magnitude that is equivalent to protective levels for SARS-CoV-2 based on clinical trial evaluation of COVID-19 vaccines (Baden et al., 2021; Polack et al., 2020). These findings suggest that repeated exposures may improve elicitation of broadly neutralizing sarbecovirus antibodies, but not broadly reactive beta-coronavirus antibodies based on the comparable cross-reactive responses to OC43 and HKU1 S observed for all cohorts. Moreover, a recent study indicated that survivors of SARS-CoV infection who subsequently received a COVID-19 vaccine had broader sarbecovirus neutralizing antibody responses than subjects only exposed to SARS-CoV-2 virus or vaccine (Tan et al., 2021). These data lend further support to the ongoing development of several vaccine candidates (Cohen et al., 2021; Martinez et al., 2021; Walls et al., 2021a) designed to specifically elicit broad sarbecovirus immunity and could protect against SARS-CoV-2 variants as well as putative future zoonotic sarbecoviruses.

## Acknowledgements

We thank Hideki Tani (University of Toyama) for providing the reagents necessary for preparing VSV pseudotyped viruses. We thank Antonio Lanzavecchia for coordination of the SARS-CoV-2 naive blood donors.

## Author contributions

A.C.W., J.E.B., and D.V. conceived the project. A.C.W. and D.V. designed the experiments and supervised this study. K.R.S., M.J.N., A.J., and C.S. carried out ELISAs. A.C.W, K.R.S, and M.J.N produced pseudovirus and ran pseudovirus neutralization assays. J.E.B., A.J., and M.M. expressed and purified proteins. E.A.G., E.J.D.A., and A.G. coordinated and ran Roche Anti N experiments. E.N.F., J.L., A.L., and H.C. coordinated collection of and provided clinical samples. A.C.W., K.R.S, M.J.N., A.J., C.S., J.E.B., and D.V. analyzed the data. A.C.W. and D.V. wrote the manuscript with input from all authors.

## Data and materials availability

Materials generated in this study will be made available on request after signing a materials transfer agreement with the University of Washington. This work is licensed under a Creative CommonsAttribution 4.0 International (CC BY 4.0) license, which permits unrestricted use, distribution, and reproduction in any medium, provided the original work is properly cited. To view a copy of this license, visit https://creativecommons.org/licenses/by/4.0/. This license does not apply tofigures/photos/artwork or other content included in the article that is credited to a third party;obtain authorization from the rights holder before using such material.

**Table S1:** Patient demographics for Delta breakthrough samples.

| Participant ID | Was this participant infected with a VOC? | Age | Sex | Is participant Hispanic or Latino | Race/Ethnicity | Which vaccine did you receive? | Days post second dose of 30 day blood draw | 30 day blood draw, days from symptom onset | Roche 30 day Anti-N reactivity (yes or no) | Days post symptom onset of 60 day draw | Days post second vaccine dose of 60 day draw | Roche 60 day Anti-N reactivity (yes or no) | Included in the 30 day group? | Included in the 60 day group? |
| --- | --- | --- | --- | --- | --- | --- | --- | --- | --- | --- | --- | --- | --- | --- |
| 267C | B.1.617.2 | 28 | Male | No | White | Pfizer | 80 | 38 | No | 63 | 105 | No | Yes | Yes |
| 268C | B.1.617.2 | 27 | Female | No | Asian | Pfizer | 94 | 28 | Yes | 65 | 131 | Yes | Yes | Yes |
| 273C | B.1.617.2 | 20 | Female | No | Asian, White | Moderna | 207 | 28 | Yes | 64 | 243 | Yes | Yes | Yes |
| 274C | S-Gene Dropout | 21 | Female | No | White | Pfizer | 194 | 26 | Yes | N/A | N/A | N/A | Yes | No |
| 275C | B.1.617.2 | 22 | Female | No | White | Pfizer | N/A | N/A | N/A | 89 | 142 | Yes | No | Yes |
| 276C | B.1.617.2 | 68 | Male | No | White | Pfizer | 212 | 24 | No | 65 | 253 | Yes | Yes | Yes |
| 277C | B.1.617.2 | 20 | Female | No | White | Pfizer | 243 | 27 | Yes | N/A | N/A | N/A | Yes | No |
| 278C | S-Gene Dropout | 26 | Female | No | White | Pfizer | 148 | 30 | Yes | 66 | 184 | Yes | Yes | Yes |
| 279C | B.1.617.2 | 27 | Female | No | White | Pfizer | 231 | 25 | Yes | N/A | N/A | N/A | Yes | No |
| 281C | B.1.617.2 | 51 | Male | No | White | Pfizer | 202 | 17 | Yes | N/A | N/A | N/A | Yes | No |
| 282C | B.1.617.2 | 21 | Female | No | Asian, Black or African American | Pfizer | 164 | 25 | Yes | N/A | N/A | N/A | Yes | No |
| 284C | S-Gene Dropout | 38 | Female | No | White | Johnson & Johnson | 166 | 20 | Yes | N/A | N/A | N/A | Yes | No |
| 285C | B.1.617.2 | 25 | Female | No | White | Moderna | 186 | 20 | Yes | 53 | 199 | Yes | Yes | Yes |
| 286C | S-Gene Dropout | 33 | Female | No | White | Pfizer | 174 | 33 | Yes | N/A | N/A | N/A | Yes | No |
| 289C | B.1.617.2 | Not Supplied | Male | No | Black or African American, White | Pfizer | N/A | N/A | N/A | 77 | 193 | Yes | No | Yes |
|  |  | Age |  |  |  |  | Days post second dose of 30 day blood draw | 30 day blood draw, days from symptom onset |  | Days post symptom onset of 60 day draw | Days post second vaccine dose of 60 day draw |  | Included in the 30 day group? | Included in the 60 day group? |
| Median: |  | 26.5 |  |  |  |  | 186 | 26 |  | 65 | 188.5 |  | n=14 | n=8 |

**Table S2:**
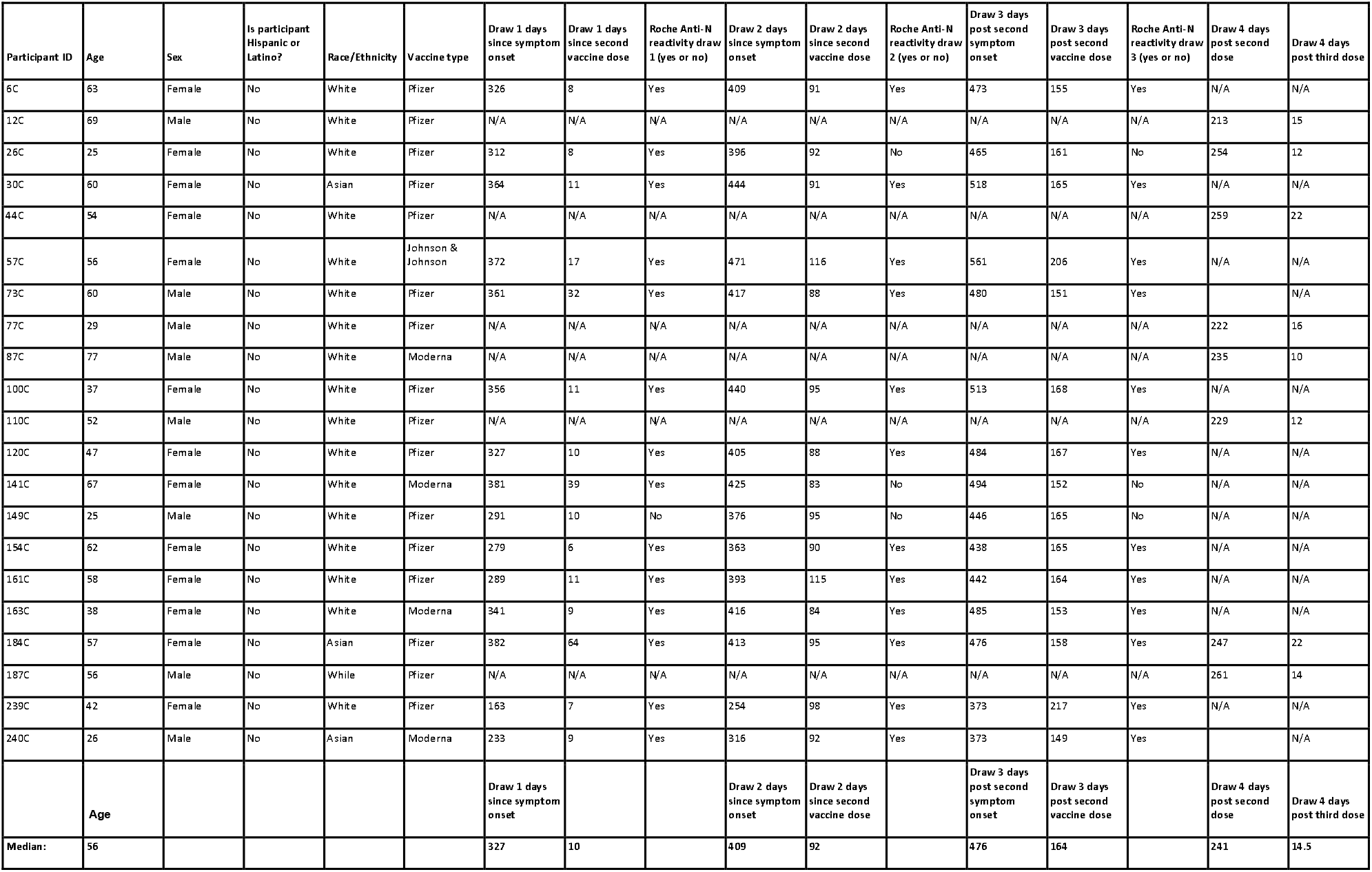
Patient demographics for SARS-CoV-2 infected/vaccinated samples.

**Table S3:**
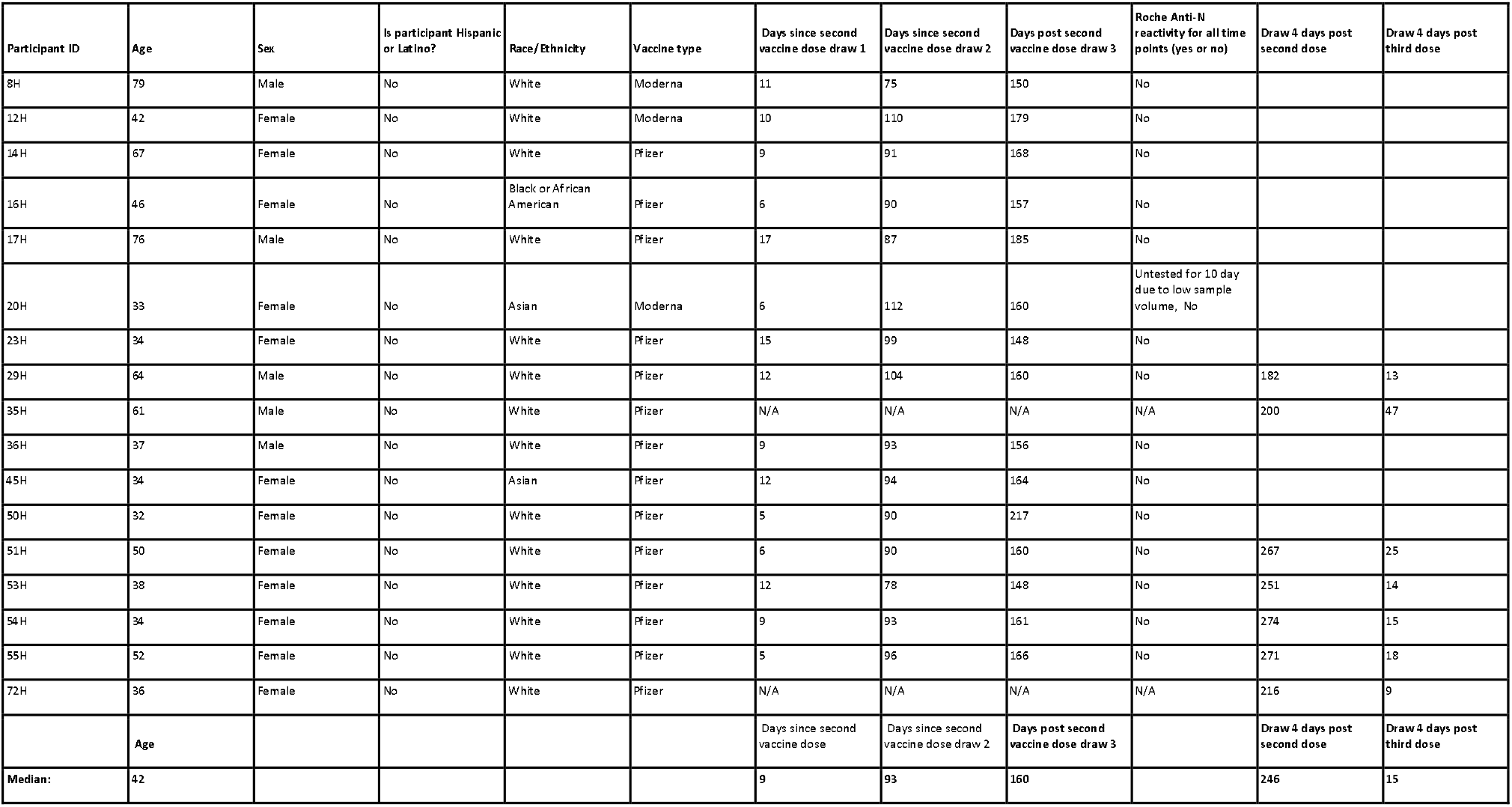
Patient demographics for vaccinated only samples.

**Table S4:**
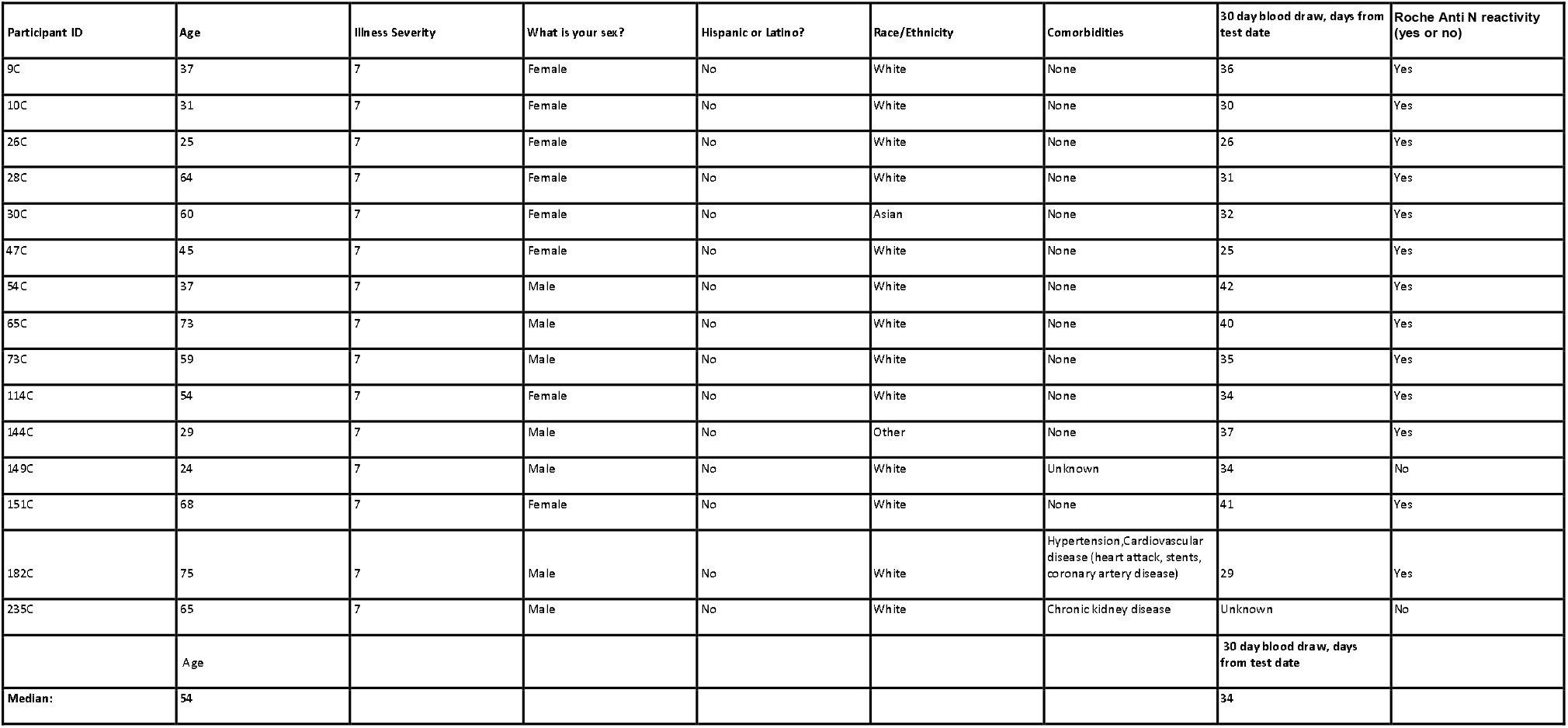
Patient demographics for SARS-CoV-2 infected only samples (HCS).

**Table S5:**
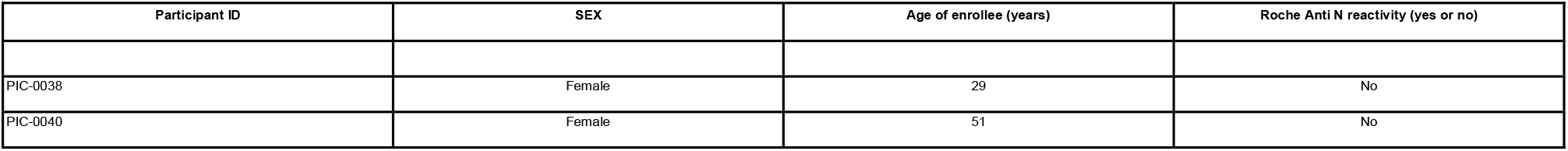
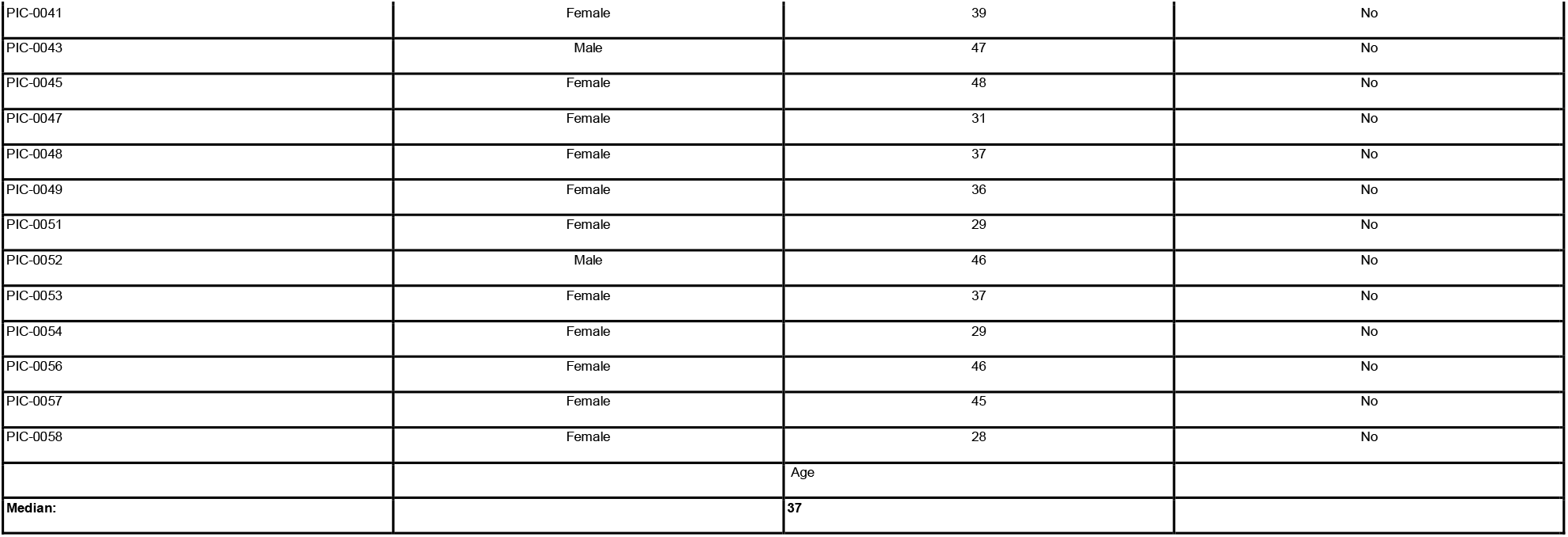
Patient demographics for SARS-CoV-2 naive samples.

**Figure S1:**
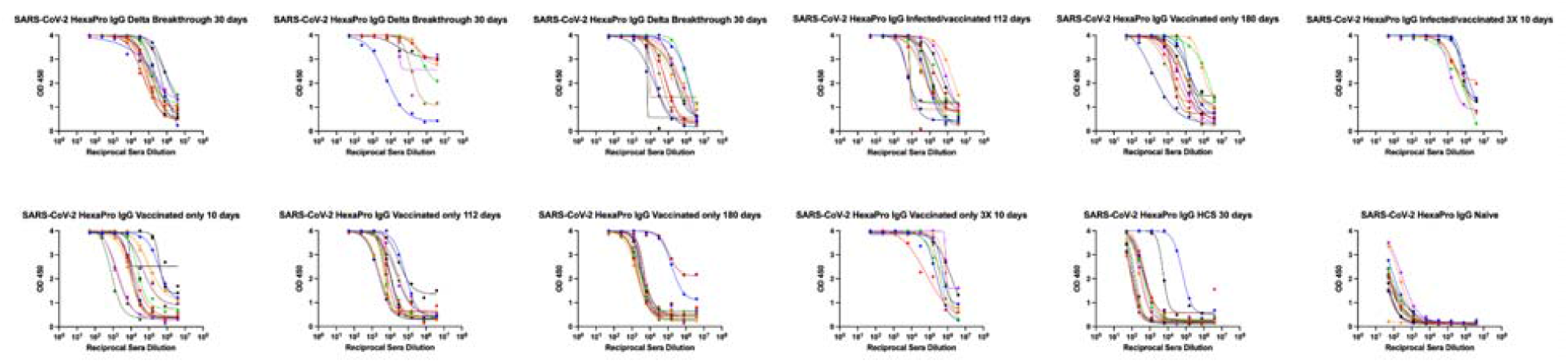
Raw SARS-CoV-2 HexaPro IgG ELISA curves and fits associated with Figure 1A, broken out by group.

**Figure S2:**
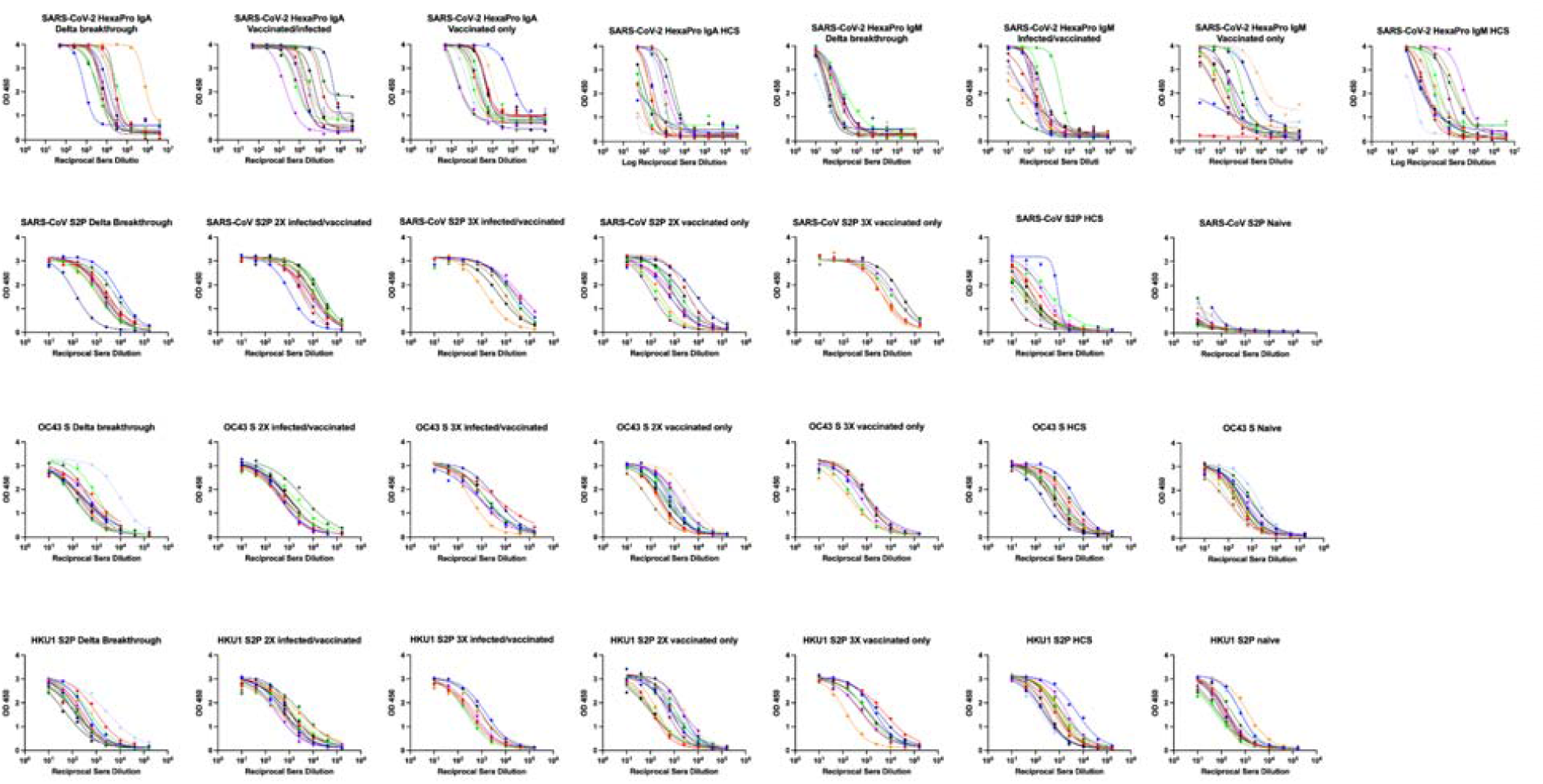
Raw ELISA curves and fits associated with figure 1B-F, broken out by group, antigen and secondary IgG. In the first row, SARS-CoV-2 HexaPro anti IgA ELISAs of Delta breakthrough, infected/vaccinated, vaccinated only, and HCS samples. In the second row, SARS-CoV-2 HexaPro anti IgM ELISAs of Delta breakthrough, infected/vaccinated, vaccinated only, and HCS samples. In the third row, SARS-CoV S 2P ELISAs of Delta breakthrough, infected/vaccinated, vaccinated only, HCS, and naive cohorts. In the fourth row, OC43 S ELISAs of Delta breakthrough, infected/vaccinated, vaccinated only, and naive cohorts. In the last row, HKU1 S2P ELISAs of Delta breakthrough, infected/vaccinated, vaccinated only, and naive samples.

**Figure S3:**
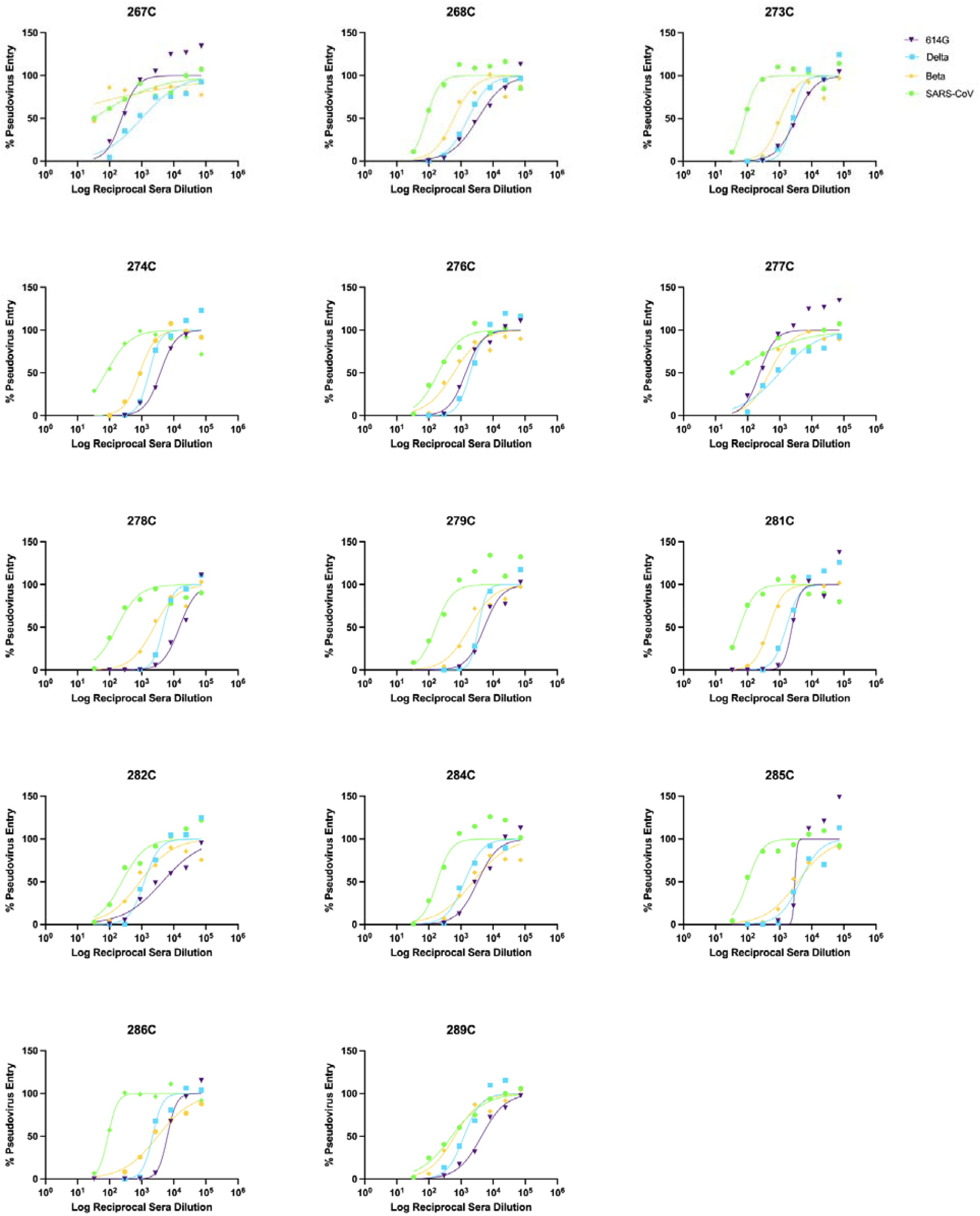
Normalized neutralization curves using VSV pseudovirus on VeroE6-TMPRSS2 cells for Delta breakthrough cases 30 days post infection.

**Figure S4:**
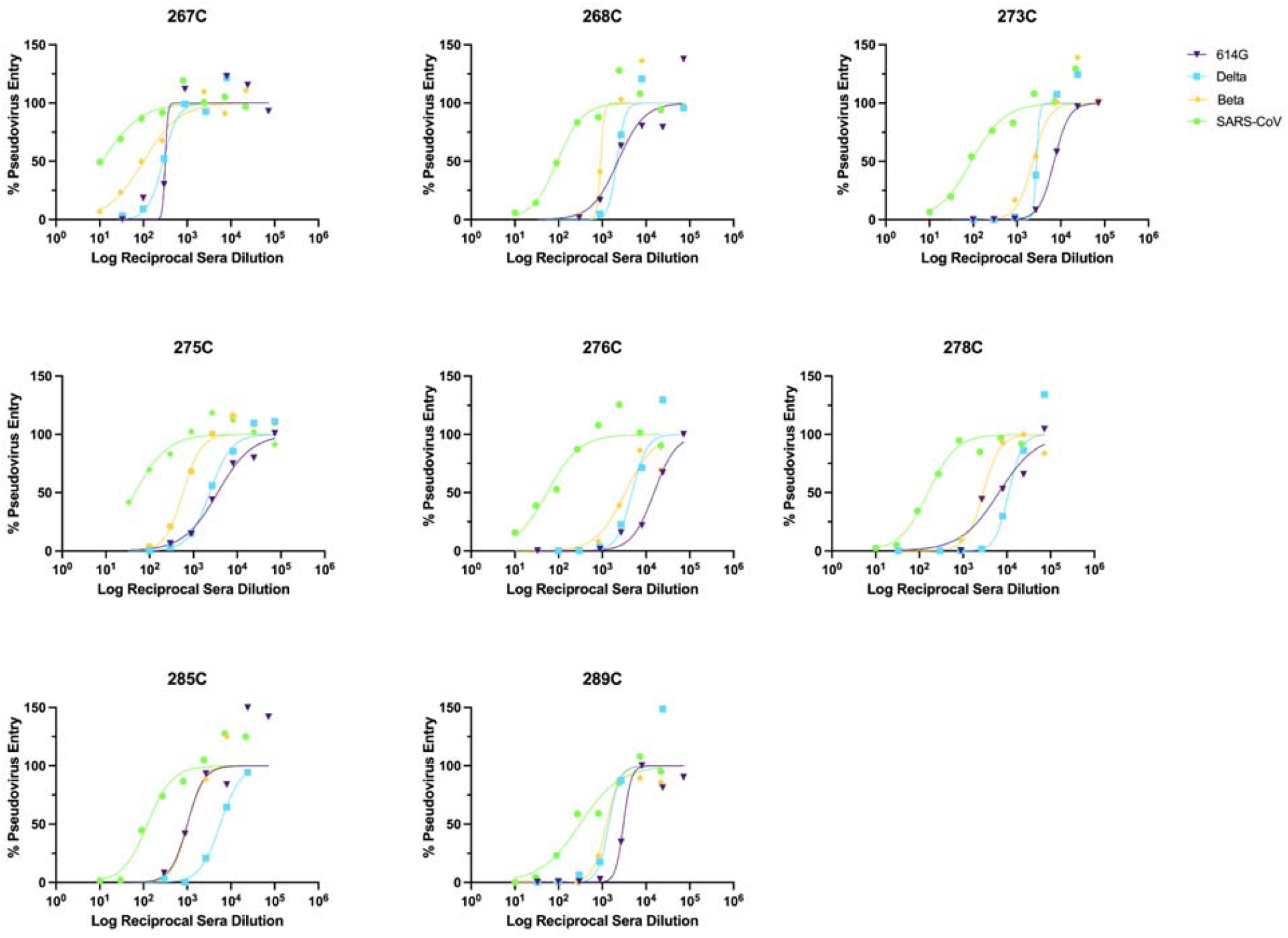
Normalized neutralization curves using VSV pseudovirus on VeroE6-TMPRSS2 cells for Delta breakthrough cases 60 days post infection.

**Figure S5:**
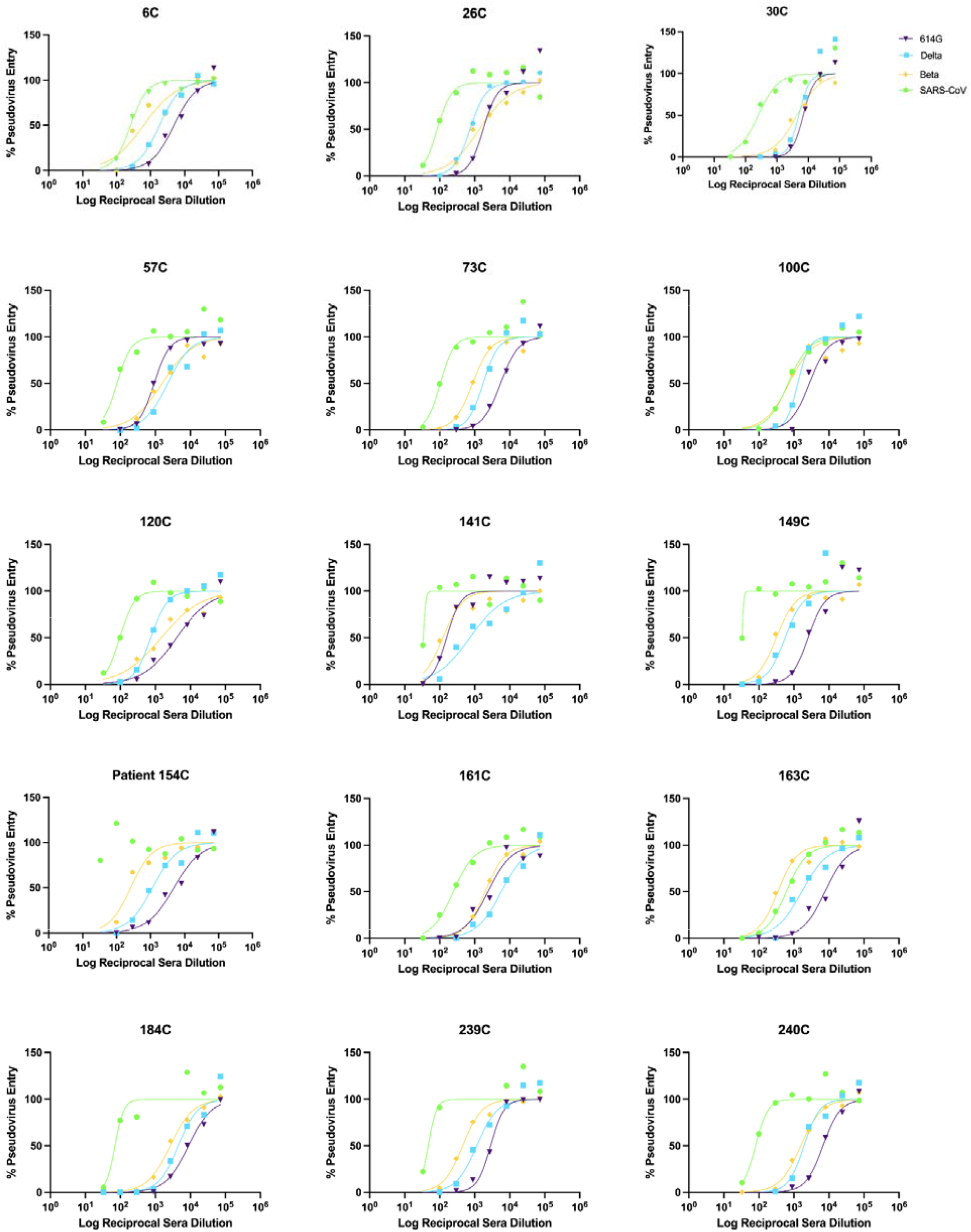
Normalized neutralization curves using VSV pseudovirus on VeroE6-TMPRSS2 cells for 2X infected/vaccinated subjects 10 days post second vaccination.

**Figure S6:**
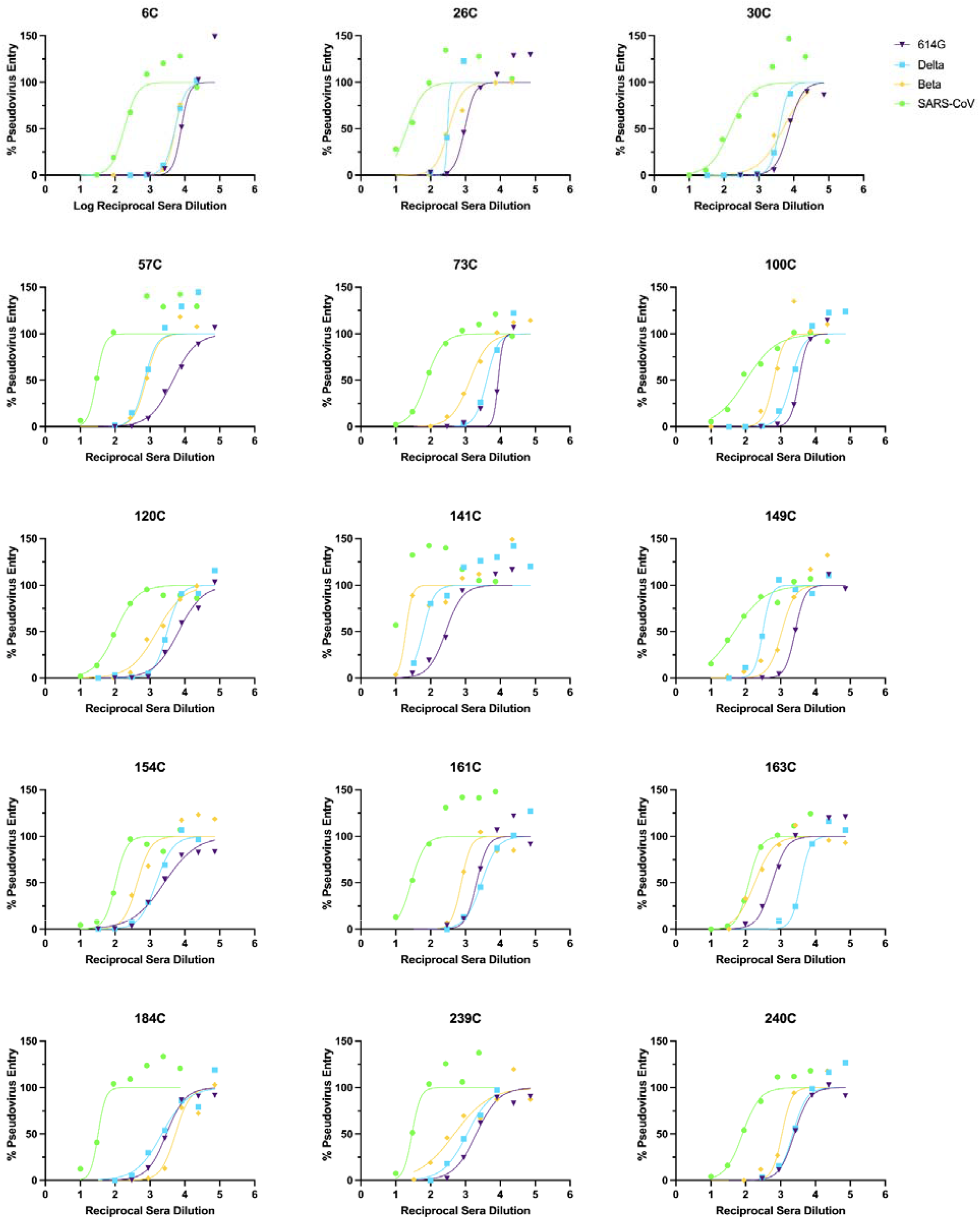
Normalized neutralization curves using VSV pseudovirus on VeroE6-TMPRSS2 cells for 2X infected/vaccinated subjects ∼112 days post second vaccination.

**Figure S7:**
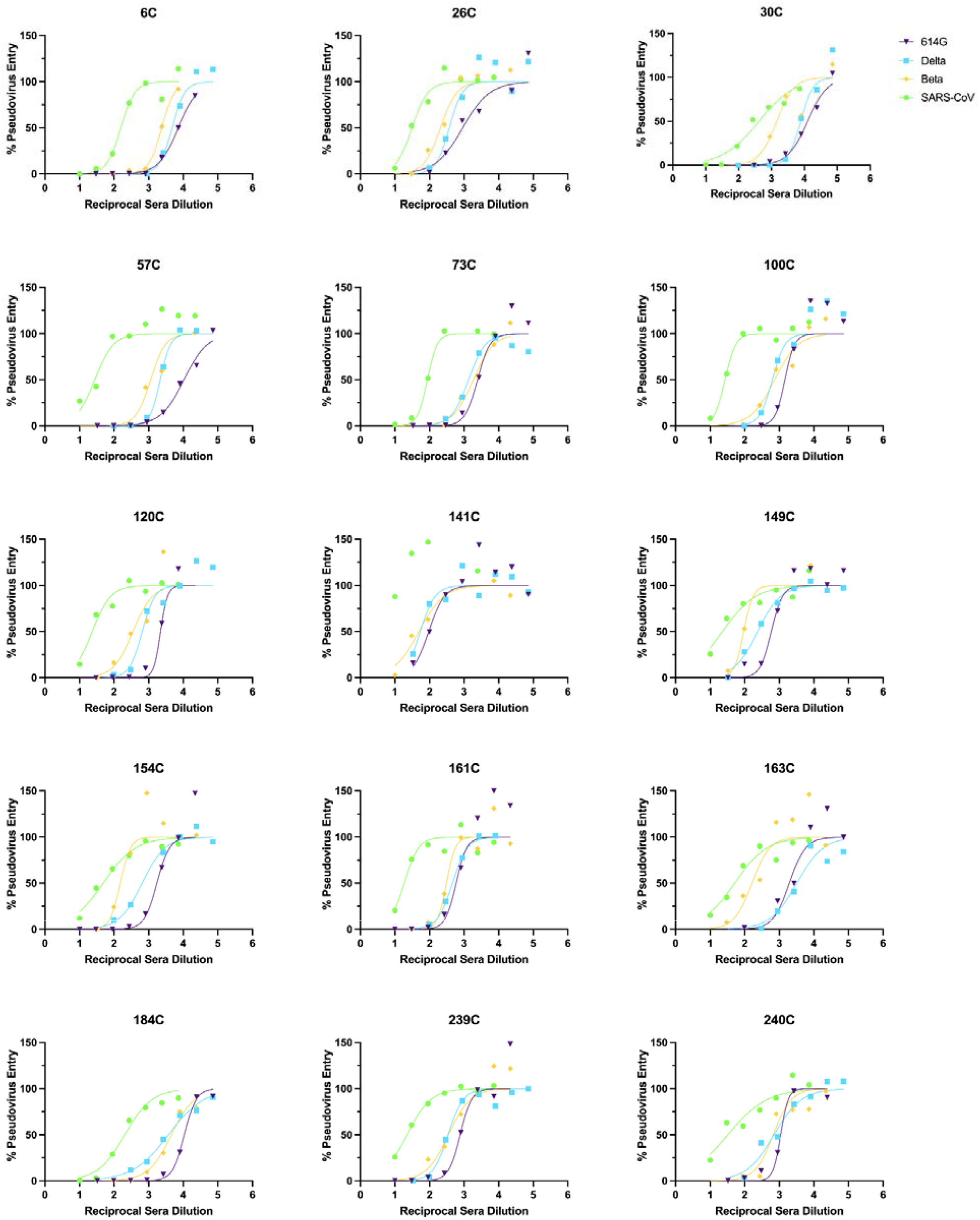
Normalized neutralization curves using VSV pseudovirus on VeroE6-TMPRSS2 cells for infected/vaccinated subjects ∼180 days post second vaccination.

**Figure S8:**
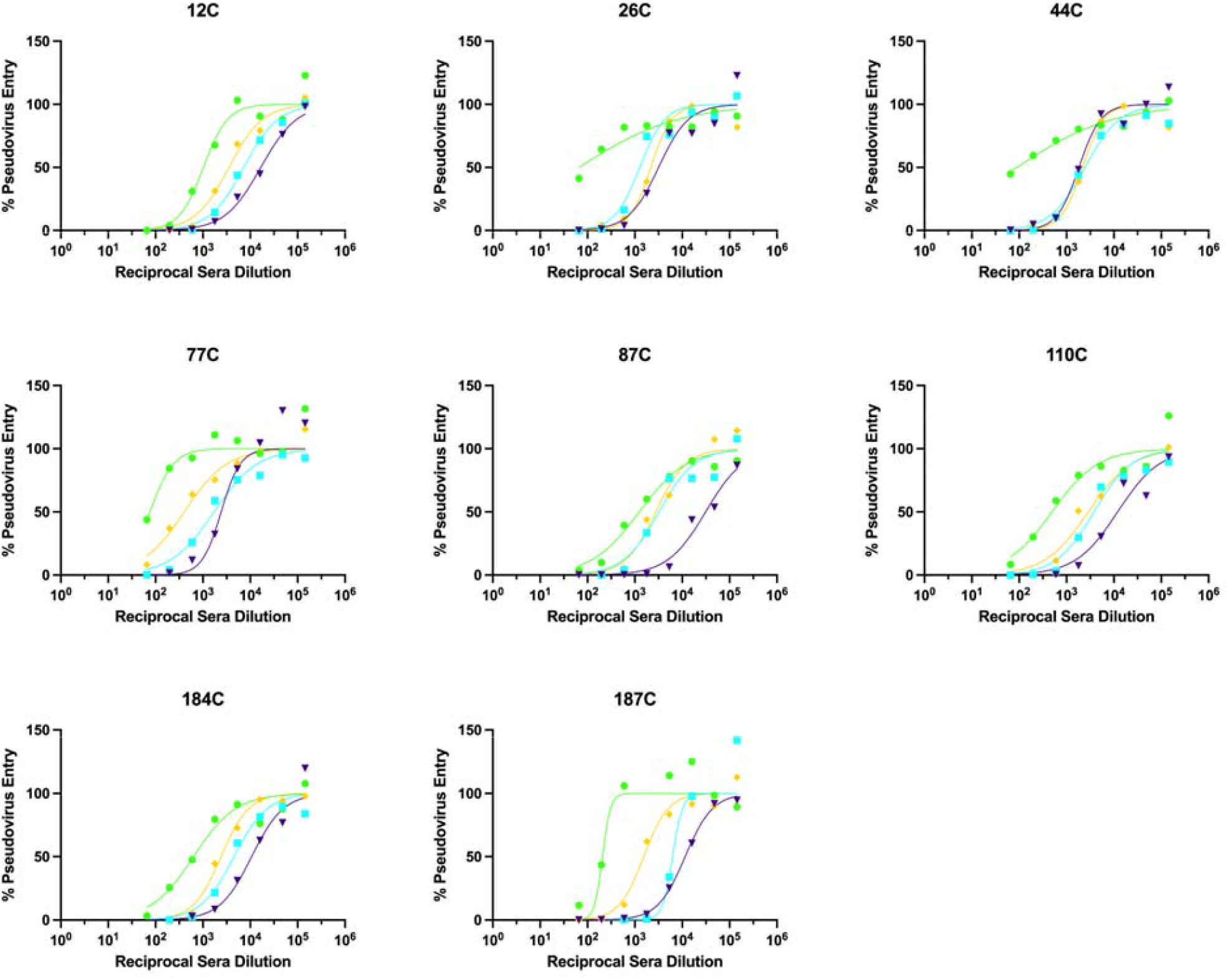
Normalized neutralization curves using VSV pseudovirus on VeroE6-TMPRSS2 cells for 3X infected/vaccinated subjects ∼10 days post third vaccination.

**Figure S9:**
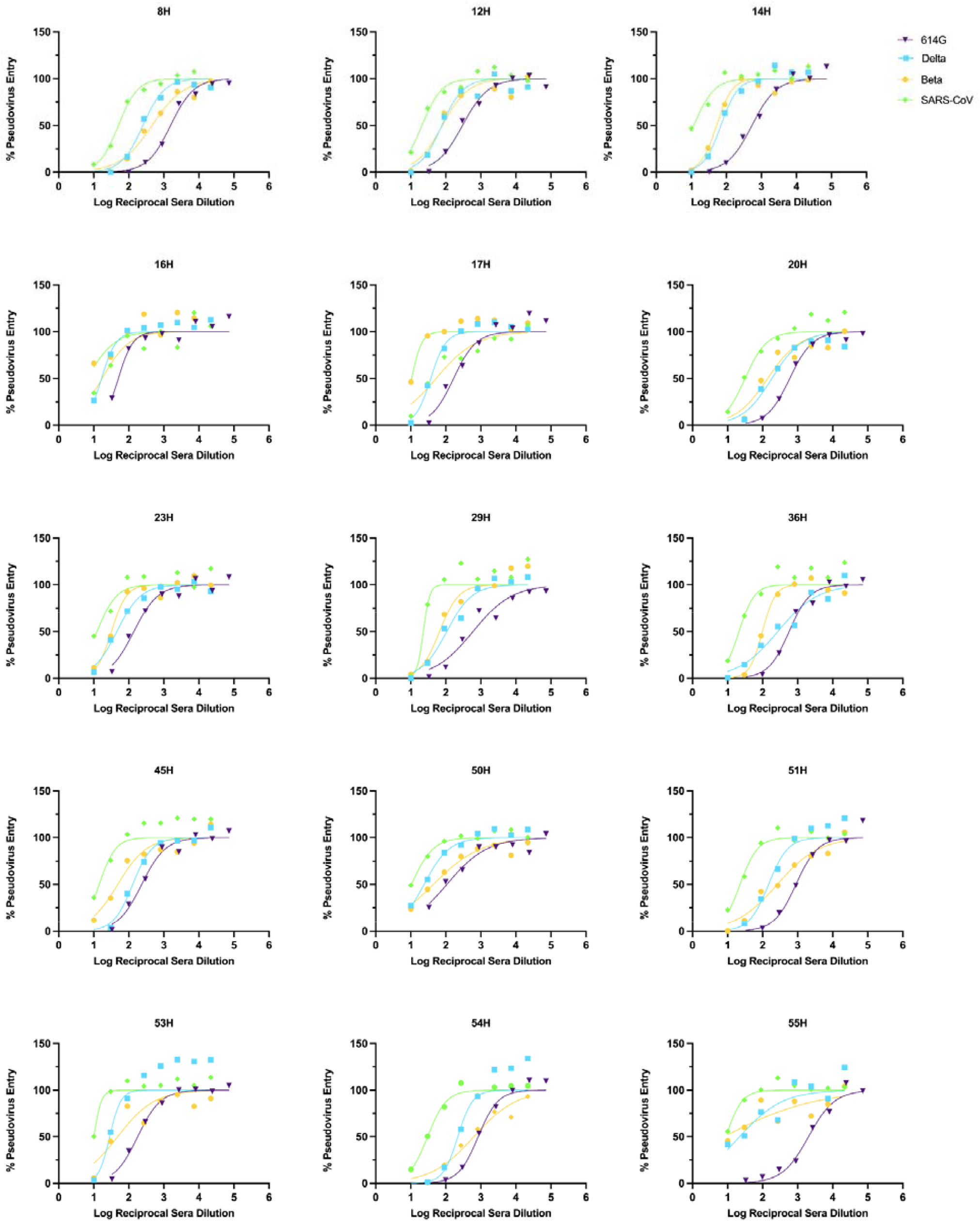
Normalized neutralization curves using VSV pseudovirus on VeroE6-TMPRSS2 cells for 2X vaccinated-only subjects 10 days post second vaccination.

**Figure S10:**
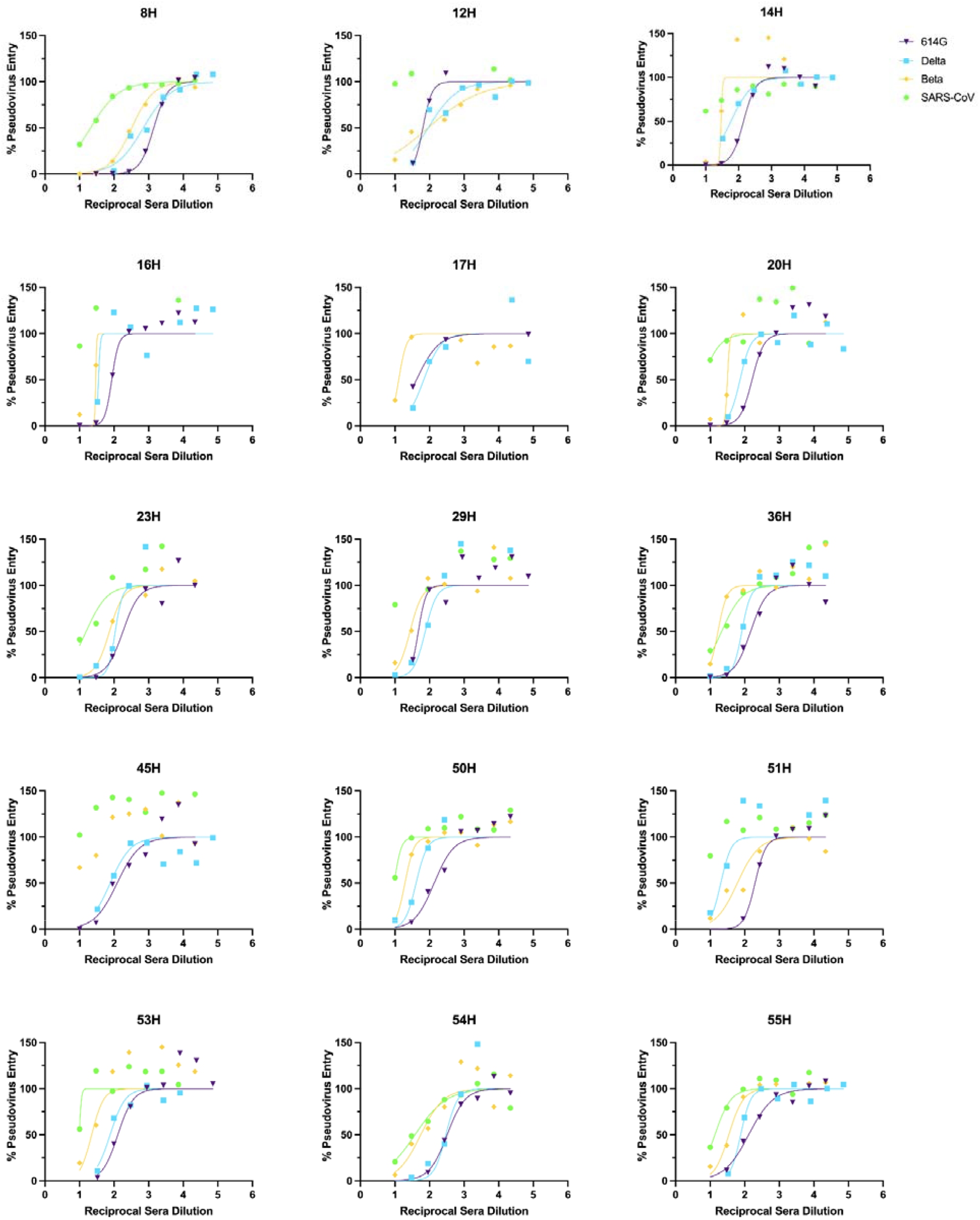
Normalized neutralization curves using VSV pseudovirus on VeroE6-TMPRSS2 cells for 2X vaccinated only subjects ∼112 days post second vaccination.

**Figure S11:**
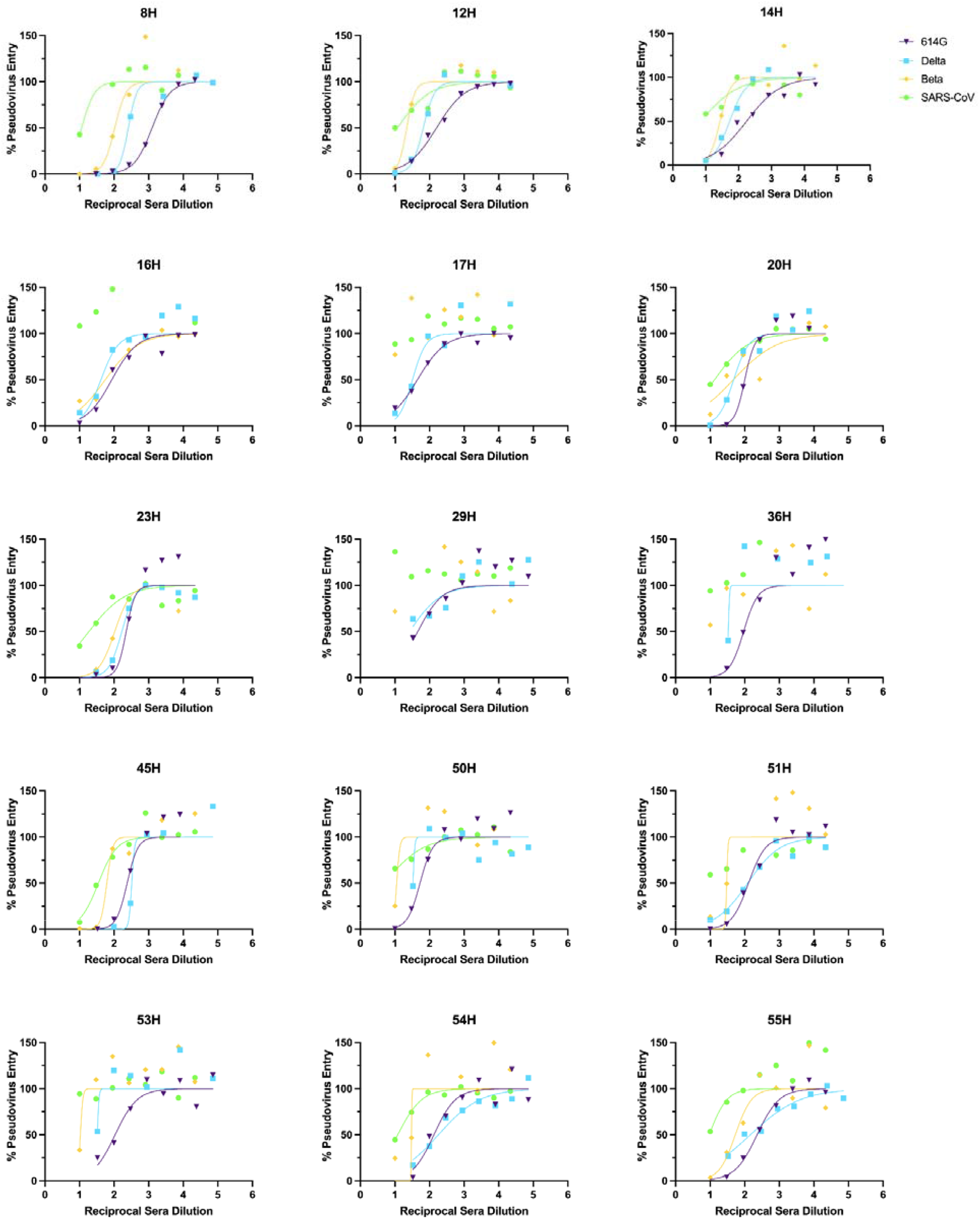
Normalized neutralization curves using VSV pseudovirus on VeroE6-TMPRSS2 cells for 2X vaccinated-only subjects ∼180 days post second vaccination.

**Figure S12:**
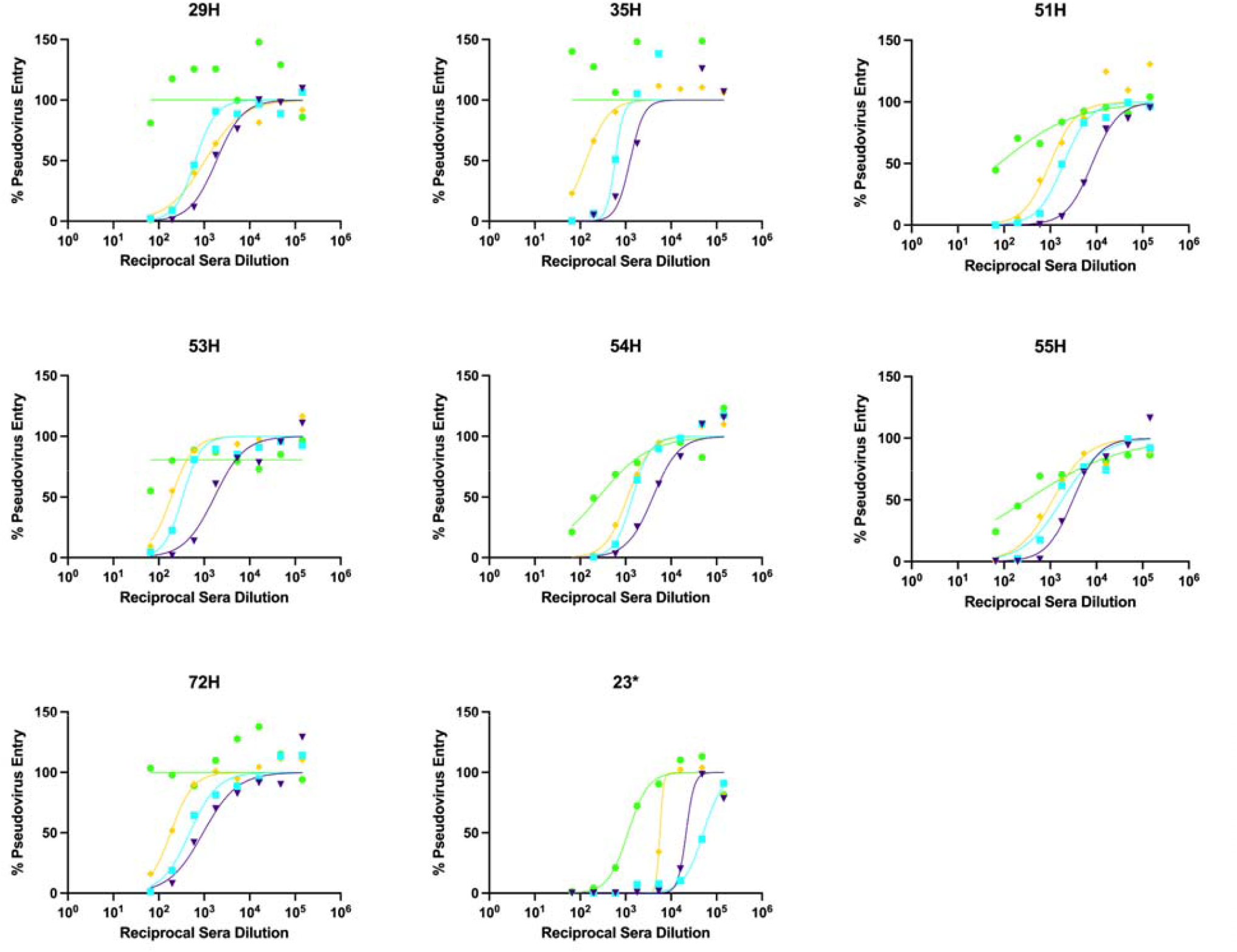
Normalized neutralization curves using VSV pseudovirus on VeroE6-TMPRSS2 cells for 3X vaccinated-only subjects ∼10 days post third vaccination.

**Figure S13:**
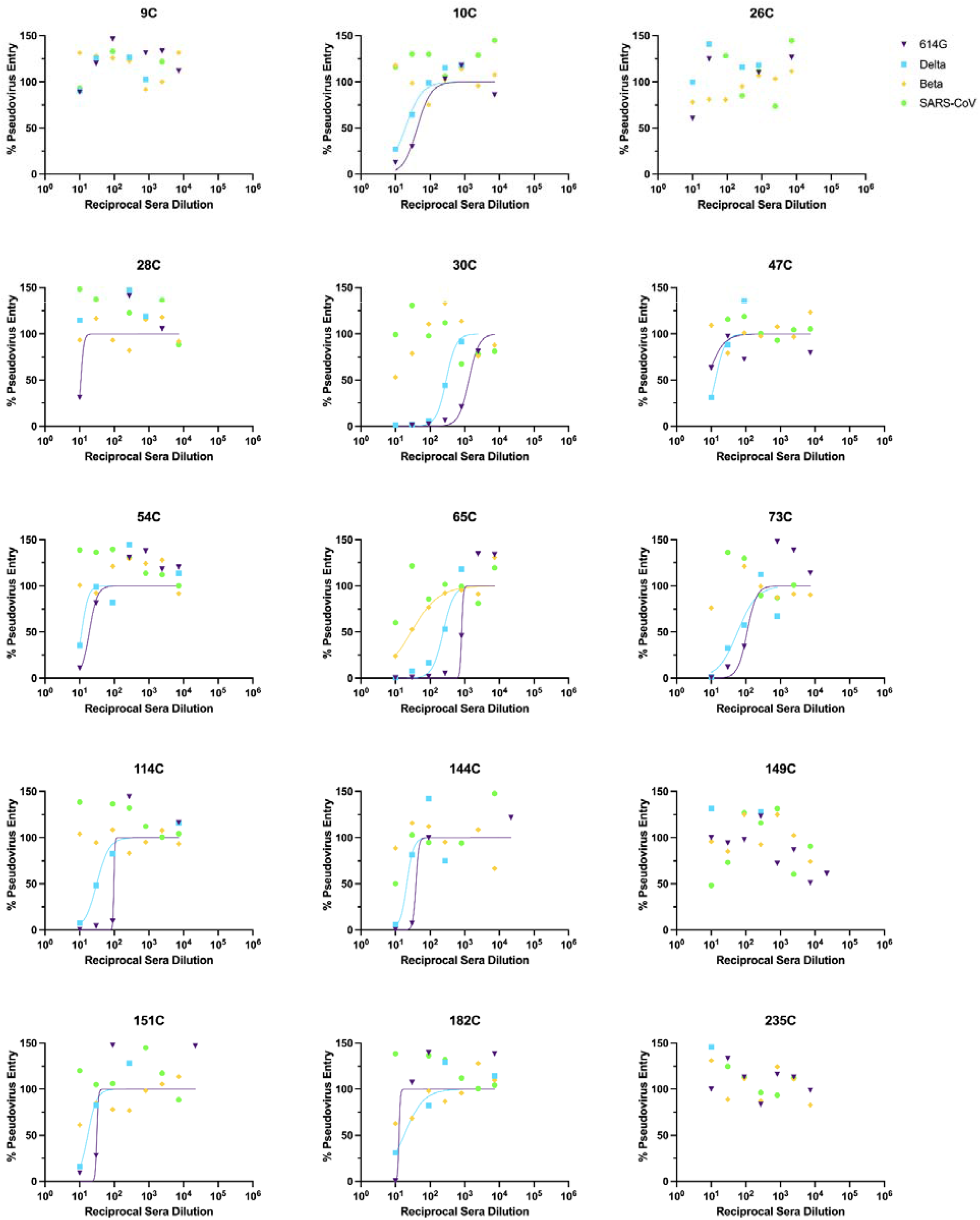
Normalized neutralization curves using VSV pseudovirus on VeroE6-TMPRSS2 cells for HCS 30 days post vaccination.

## Methods

### Human Sera/Plasma

Blood samples were collected from participants as part of the Hospitalized or Ambulatory Adults with Respiratory Viral Infections (HAARVI) study and was approved by the University of Washington Human Subjects Division Institutional Review Board (STUDY00000959). Baseline socio-demographic and clinical data for these individuals are summarized in Tables S1-S5. All samples were sera except for the 60 day Delta breakthrough samples which were ACD plasma.

### Sequencing of Delta Breakthrough Individuals

Participants were enrolled after SARS-CoV-2 positive RT-PCR results with S-Gene Dropout. SARS-CoV-2 genome sequencing was then conducted using a targeted enrichment approach to confirm Delta variant status. RNA from positive specimens was converted to cDNA using random hexamers and reverse transcriptase (Superscript IV, Thermo) and a sequencing library was constructed using the Illumina TruSeq RNA Library Prep for Enrichment kit (Illumina). The sequencing library was enriched for SARS-CoV-2 using the Twist Respiratory Virus Research Panel (Twist). Libraries were sequenced on a MiSeq or NextSeq instruments. The resulting reads were assembled against the SARS-CoV-2 reference genome Wuhan-Hu-1/2019 (Genbank accession MN908947) using the bioinformatics pipeline https://github.com/seattleflu/assembly. Consensus sequences were deposited to Genbank and GISAID (Bedford et al., 2020). Sequence quality was determined using Nextclade version 1.0.0-alpha.8 (https://clades.nextstrain.org/). Lineage was assigned using the Pangolin COVID-19 Lineage Assigner version 3.1.11 (https://pangolin.cog-uk.io/)(Paredes et al., 2021).

### Recombinant Protein Expression and purification

The SARS-CoV-2 Hexapro S (Hsieh et al., 2020), SARS-CoV-2 Delta VFLIP S with the native furin cleavage site (McCallum et al., 2021a; Olmedillas et al., 2021), SARS-CoV S 2P (Pallesen et al., 2017; Walls et al., 2020b), HKU1 S2P, and OC43 S(Tortorici et al., 2019) ectodomains were produced in Expi293F Cells (ThermoFisher Scientific) grown in suspension using Expi293 Expression Medium (ThermoFisher Scientific) at 37°C in a humidified 8% CO2 incubator with constant rotation at 130 rpm. Cells grown to a density of 3 million cells per mL were transfected using the ExpiFectamine 293 Transfection Kit (ThermoFisher Scientific) and cultivated for four days at which point the supernatant was harvested. SARS-CoV-2 Hexapro S, SARS-CoV-2 Delta VFLIP S, SARS-CoV S2P, and HKU1 S2P ectodomains were purified from clarified supernatants using a HisTrapHP column (Cytiva) and washed with 10 column volumes of 25 mM sodium phosphate pH 8.0 and 150 mM NaCl before elution on a gradient up to 500 mM imidazole. OC43 S fused to strep tag, the supernatant was clarified by 10 minute centrifugation at 800xg and brought to 100 mM Tris-HCL pH 8, 150 mM NaCl, and 18.1 mL/liter biotin blocking solution (BioLock). After a 20-minute incubation, supernatant was further centrifuged at 10,000xg for 20 minutes. Supernatant was then bound to a 1 mL StrepTrap HP column (Cytiva) and washed with 10 column volumes of 100 mM Tris-HCl, 150 mM NaCl, and 1 mM EDTA pH 8 before elution in wash buffer supplemented with 2.5 mM desthiobiotin. Purified protein was buffer exchanged into 20 mM Tris-HCl pH 8.0 and 100 mM NaCl, concentrated using 100 kDa MWCO centrifugal filters (Amicon Ultra) to 1-2 mg/mL, and flash frozen.

### ELISA

For anti-S ELISA, 50 μL of 2-6 μg/mL S was plated onto 384-well Nunc Maxisorp (ThermoFisher) plates in PBS and sealed overnight at 4°C. The next day plates were washed 4 × in Tris Buffered Saline Tween (TBST-20mM Tris pH 8, 150mM NaCl, 0.1% Tween) using a plate washer (BioTek) and blocked with Casein (ThermoFisher) for 1 h at 37°C. Plates were washed 4 × in TBST and 1:5 serial dilutions of human sera or plasma were made in 50 μL TBST starting at 1:10, 1:50, or 1:250 and incubated at 37°C for 1 h. Plates were washed 4 × in TBST, then anti-human (Invitrogen) horseradish peroxidase-conjugated antibodies were diluted 1:5,000 and 50 μL added to each well and incubated at 37°C for 1 h. Plates were washed 4 × in TBST and 50 μL of TMB (SeraCare) was added to every well for 5 min at room temperature. The reaction was quenched with the addition of 50 μL of 1 N HCl. Plates were immediately read at 450 nm on a VarioSkanLux plate reader (ThermoFisher) and data plotted and fit in Prism (GraphPad) using nonlinear regression sigmoidal, 4PL, X is log(concentration) to determine EC_50_ values from curve fits. Where the curve did not reach an OD450 of 4, a constraint of OD450 4 was placed on the upper bounds of the fit.

### VSV Pseudovirus Production

G614 SARS-CoV-2 S (YP 009724390.1), Delta S, Beta S, and SARS-CoV S pseudotyped VSV viruses were prepared as described previously (McCallum et al., 2021a; Walls et al., 2021b). Briefly, HEK293T cells in DMEM supplemented with 10% FBS, 1% PenStrep seeded in 10-cm dishes were transfected with the plasmid encoding for the corresponding S glycoprotein using lipofectamine 2000 (Life Technologies) following the manufacturer’s instructions. One day post-transfection, cells were infected with VSV(G*ΔG-luciferase)(Kaname et al., 2010) and after 2 h were washed five times with DMEM before adding medium supplemented with anti-VSV-G antibody (I1-mouse hybridoma supernatant, CRL-2700, ATCC). Virus pseudotypes were harvested 18-24 h post-inoculation, clarified by centrifugation at 2,500 x g for 5 min, filtered through a 0.45 μm cut off membrane, concentrated 10 times with a 30 kDa cut off membrane, aliquoted and stored at −80°C.

### VSV Pseudovirus Neutralization

VeroE6-TMPRSS2 (Lempp et al., 2021) were cultured in DMEM with 10% FBS (Hyclone), 1% PenStrep and 8 μg/mL puromycin (to ensure retention of TMPRSS2) with 5% CO_2_ in a 37°C incubator (ThermoFisher). Cells were trypsinized using 0.05% trypsin and plated to 40,000 cells/well. The following day, cells were checked to be at 80% confluence. In an empty half-area 96-well plate, a 1:3 serial dilution of sera was made in DMEM and diluted pseudovirus was then added and incubated at room temperature for 30-60 min before addition of the sera-virus mixture to the cells at 37°C. 2 hours later, 40 μL of a DMEM solution containing 20% FBS and 2% PenStrep was added to each well. After 17-20 hours, 40 μL/well of One-Glo-EX substrate (Promega) was added to the cells and incubated in the dark for 5-10 min prior to reading on a BioTek plate reader. Measurements were done at least in duplicate using distinct batches of pseudoviruses and one representative experiment is shown. Relative luciferase units were plotted and normalized in Prism (GraphPad). Nonlinear regression of log(inhibitor) versus normalized response was used to determine IC_50_ values from curve fits. Normality was tested using the D’agostino-Pearson test and in the absence of a normal distribution, kruskal-wallis tests were used to compare two groups to determine whether differences reached statistical significance. Fold changes were determined by comparing individual animal IC_50_ and then averaging the individual fold changes for reporting.

